# Two conserved arginine residues facilitate C-S bond cleavage and persulfide transfer in Suf family cysteine desulfurases

**DOI:** 10.1101/2024.10.17.618868

**Authors:** Rajleen K. Gogar, Juliana V. Conte, Jack A. Dunkle, Patrick A. Frantom

**Author notes:** to whom correspondence should be addressed: Patrick A. Frantom,; Jack A. Dunkle.

## Abstract

Under conditions of oxidative stress or iron starvation, iron-sulfur cluster biogenesis in *E. coli* is initiated by the cysteine desulfurase, SufS, via the SUF pathway. SufS is a type II cysteine desulfurase that catalyzes the PLP-dependent breakage of an L-cysteine C-S bond to generate L-alanine and a covalent active site persulfide as products. The persulfide is transferred from SufS to SufE and then to the SufBC_2_D complex, which utilizes it in iron-sulfur cluster biogenesis. Several lines of evidence suggest two conserved arginine residues that line the solvent side of the SufS active site could be important for function. To investigate the mechanistic roles of R56 and R359, the residues were substituted using site-directed mutagenesis to obtain R56A/K and R359A/K SufS variants. Steady state kinetics indicated R56 and R359 have moderate defects in the desulfurase half reaction but major defects in the transpersulfurase step. Fluorescence polarization binding assays showed that the loss of activity was not due to a defect in forming the SufS/SufE complex. Structural characterization of R56A SufS shows loss of electron density for the α3-α4 loop at the R56/G57 positions, consistent with a requirement of R56 for proper loop conformation. The structure of R359A SufS exhibits a conformational change in the α3-α4 loop allowing R56 to enter the active site and mimics the residue’s position in the PLP-cysteine aldimine structure. Taken together, the kinetic, binding, and structural data support a mechanism where R359 plays a role in linking SufS catalysis with modulation of the α3-α4 loop to promote a close-approach interaction of SufS and SufE conducive to persulfide transfer.

## Introduction

Sulfur is an essential element for life and is involved in the formation of a variety of biomolecules such as amino acids, coenzymes, modified tRNA bases, and iron-sulfur clusters. Mobilization of sulfur often involves cysteine desulfurases that use a pyridoxal 5’-phosphate (PLP) cofactor to catalyze the breakage of the L-cysteine C-S bond yielding L-alanine and covalent persulfide on an active site cysteine (Cys-S-S^−^) as products.(1, 2) The model organism *Escherichia coli* contains three cysteine desulfurases in its genome, two of which participate in bioassembly pathways for Fe-S cluster cofactors. IscS catalyzes the cysteine desulfurase step in the housekeeping ISC pathway passing the nascent persulfide intermediate directly to IscU for cluster formation under normal coditions.(3) IscS is capable of interacting with multiple persulfide acceptors and also supports formation of additional thio-containing metabolites.(4) Under conditions of oxidative stress or iron-limitation, *E. coli* utilizes the SUF pathway as an alternative to the ISC pathway (Figure 1A).(5, 6) The SUF pathway has been shown to be resistant to external oxidants and reductants, consistant with its activation under harsh conditions.(7, 8) The sulfur mobilization step in the SUF pathway is catalyzed by the cysteine desulfurase SufS (Figure 1B). Due to this protective function, SufS requires a specific transpersulfurase partner, SufE, to act as an intermediary and transfer the persulfide to the SufBC_2_D scaffold for assembly into iron-sulfur clusters.(9, 10) In contrast to IscS, persulfide trafficking through the SUF pathway is dedicated to Fe-S cluster assembly.

**Figure 1.**
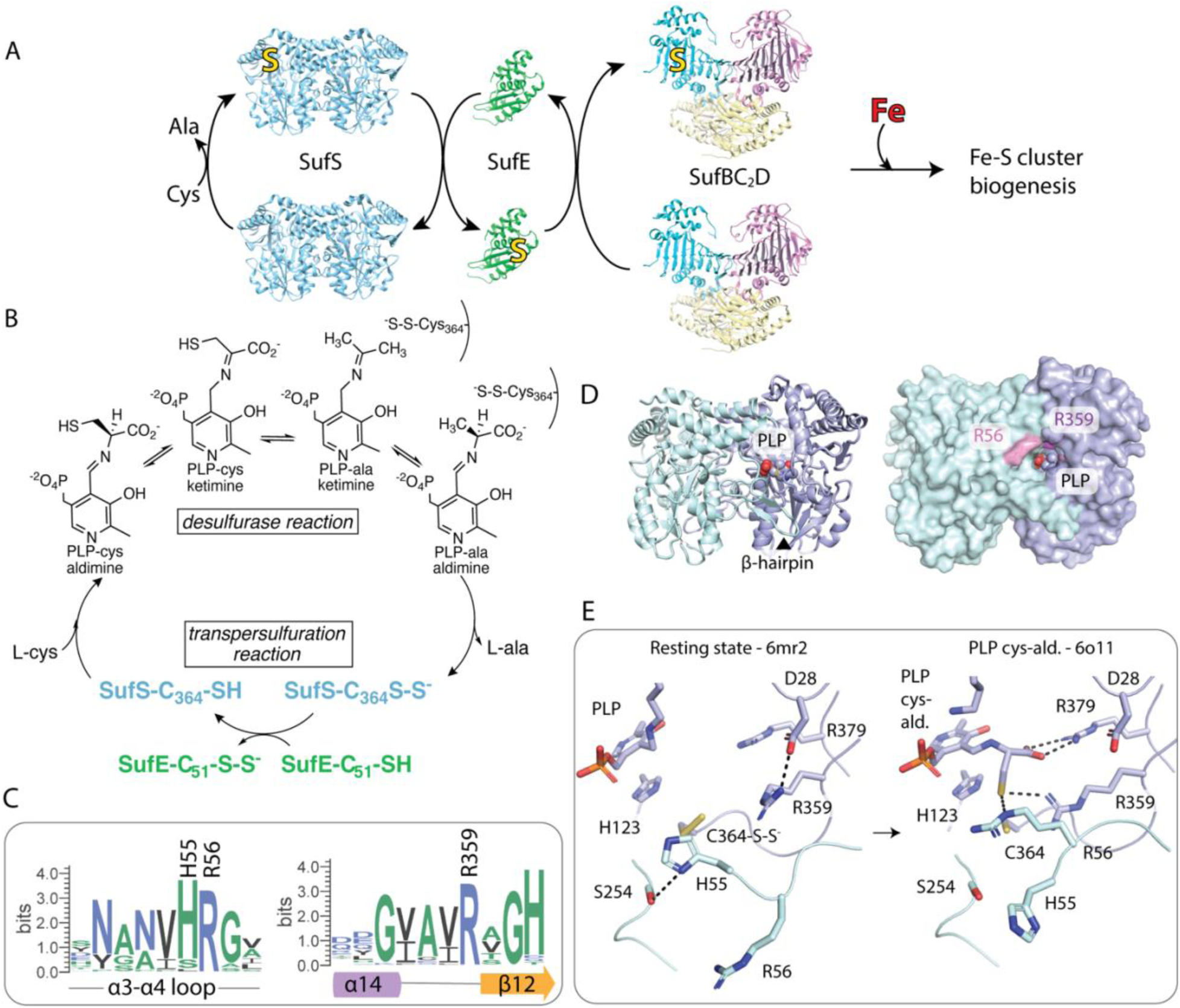
The location of conserved Arg residues in the SufS active site. (A) Sulfur liberated from Cys by SufS is utilized by the Suf pathway for iron-sulfur cluster biogenesis. (B) SufS catalyzes a desulfurase reaction to generate a covalently bound persulfide which is then transferred to SufE during the transpersulfuration reaction. Several covalent intermediates form during the desulfurase reaction. (C) Sequence logos display the high level of conservation for R56 and R359 in type II cysteine desulfurases. (D) SufS has a homodimeric structure and its active site is marked by the PLP cofactor. A β-hairpin structure lies adjacent to the active site. A surface rendering of SufS shows that R56 (pink) and R359 (purple) line the channel to the active site. (E) SufS crystal structures bearing the PLP cys-aldimine reaction intermediate reveal that R56 and R359 undergo conformational changes to form interactions with the substrate at this step. The structures shown are based on PDB codes 6mr2, 6o11. PDB code 6o11 contains an active site mutant that blocks chemistry but is rendered here as wild-type to mimic the natural state.

Both SufS and IscS belong to the family of homodimeric aminotransferase type V-fold PLP-dependent enzymes; however, there are several key structural differences that separate the two enzymes. IscS is an example of a type I cysteine desulfurase as its active site cysteine residue is located on a long, flexible loop.(11) The flexible loop structure allows IscS to directly transfer the persulfide product to the IscU scaffold for cluster assembly.(12) SufS is a type II cysteine desulfurase with its active site cysteine located on a short, rigid loop buried ∼10 Å below the active site entrance(13), necessitating the requirement of SufE in the SUF pathway. SufS also contains a sequence insertion relative to the type I enzymes that forms a β-hairpin that partially covers the active site of the adjacent monomer (Figure 1D). This β-hairpin is part of a larger β-latch motif responsible for protection of the persulfide and formation of a “close-approach” complex with SufE.(14, 15)

An abbreviated chemical mechanism for SufS is shown in Figure 1B. SufS is isolated with a PLP cofactor attached to K226 (*E. coli* numbering) as an internal aldimine. In the presence of L-cysteine, a transamination reaction follows to give an external PLP-Cys aldimine. The structure of this enzyme intermediate was trapped using the C364A variant of SufS and characterized by X-ray crystallography (PDB: 6o11).(16) A comparison of the PLP-Cys aldimine intermediate with the internal aldimine SufS structure or “resting state” (PDB: 6mr2) identifies three active site residues that change conformation, H55, R56 and R359, all of which are strongly conserved in type II cysteine desulfurases (Figure 1C). In the 6mr2 structure, which contains a persulfide intermediate on C364, H55 is found pointing into the active site and interacts with the sidechain of S254, while R56 and R359 face out of the active site (Figure 1E). S254 is located in the β-latch structural element that helps form a high affinity “close-approach” complex with SufE.(14) H55 and R56 lie on the α3-α4 loop (residues 50-59), which has also recently been implicated as important for SufS/SufE interactions.(17) R359 lies on a β-strand adjacent to the active site and makes contact with the sidechain of D28 as well as residues on the flexible H59 sidechain and H55 carbonyl oxygen. We define the positions of R56 and R359 as “out” in this structure.

These three residues shift conformation in the 6o11 structure containing the PLP-Cys aldimine intermediate and make interactions with the sulfur atom on the aldimine (Figure 1E). In this “in” position, R359 is poised to interact with both the PLP-Cys external aldmine sulfur and the C364 sidechain suggesting it may play a key role in chemistry. The α3-α4 loop undergoes a conformational change allowing the terminal group of R56 to swing ∼ 10 Å into the active site (“in” position) and H55 to swing out. In the active site, R56 makes contact with the sulfur of the PLP-Cys aldimine as well as T278 and the backbone carbonyls of G277 and V54. Based on this series of structures and proposed roles for the α3-α4 loop and the β-latch structural element, we hypothesize that coordinated motions of R359 and the α3-α4 loop work together to link the catalytic mechanism of SufS with protected persulfide transfer to SufE.

To investigate this hypothesis, site-directed mutants of conserved R359 and R56 residues in SufS were generated and characterized by cysteine desulfurase kinetics, SufS/SufE binding studies, and X-ray crystallography. The H55A SufS variant has been previously characterized and shown to have minimal effects on catalysis and structure.(16) In contrast, substitution of either the R56 and R359 residue with alanine results in catalytically impaired SufS variants, with the partial recovery of enzyme activity upon substitution with lysine. Fluorescence polarization binding assays show catalytic defects in the alanine variants are not due to disruption of the SufS/SufE complex. X-ray crystal structures of R56A and R359A SufS variants show structural changes in the active site focused on the conformation of the α3-α4 loop. Importantly, the R359A SufS structure shows an α3-α4 loop conformation mimicking the PLP-Cys aldimine structure, which could promote SufE interactions. Overall, the data suggest an orchestrated mechanism followed by R56 and R359 SufS in response to the changes in the active site, where the positioning of the R359 SufS residue governs the movement of the α3-α4 mobile loop and in turn interactions with SufE.

## Experimental Procedures

### Generation of site-directed SufS variants

Mutations to generate the SufS variants R56A, R56K, R359A, R359K were introduced into the wildtype pET-21a-*sufS* vector via Quick Change Site-Directed Mutagenesis Kit (Agilent) using forward and reverse primers with desired mutations (Table S1). The resulting plasmids were transformed into XL-10 Gold *E. coli* cells, purified, and sequenced to confirm the desired mutations (Eurofin Genomics). Recombinant protein expression for SufS, SufE, and SufS variants was done by transforming pET-21a-*sufS* plasmids carrying the desired mutations into electrocompetent BL21(DE3)*Δsuf E. coli* cells, which lack the genomic *suf* operon. Next, cells were grown overnight in Luria Broth (LB) with 100 μg/ml ampicillin and 50 μg/ml kanamycin at 37 °C. The next morning the overnight cultures were diluted 100-fold in fresh LB containing 100 μg/ml ampicillin and 50 μg/ml kanamycin. The cells were incubated at 37 °C with shaking at 250 rpm until an A_600_ of 0.4-0.6 was reached. At this point, protein expression was induced with 500 μM IPTG. Cells were grown for 3 h post induction and harvested by centrifugation. Cell pellets were stored at −80 °C until purification. For SufS purification, cells were thawed and resuspended in lysis buffer consisting of 25 mM Tris pH 7.5, 100 mM NaCl, 5 mM DTT, 0.02 mg/mL DNase, 1 mM PMSF, and 1 mM MgCl_2_. Resuspended cells were lysed via sonication for 10 min (30 sec on and 30 sec off) at output control of 4 and duty cycle of 50% on a Branson sonifier. The lysate was centrifuged at 20,000g for 30 min at 4 °C to pellet cell debris. SufS and SufE proteins were purified using methods described previously.(18) Briefly, WT SufS and SufS mutants were purified using three sets of columns in a sequence of Q-XL anion-exchange, phenyl-HP hydrophobic interactions, and Superdex-200 size exclusion chromatography (Cytivia). Lysate containing wildtype SufE was purified using two sets of columns in a sequence of Q-XL anion exchange and Superdex-200 size exclusion chromatography. Fractions containing desired proteins (SufS or SufE) were identified using SDS-PAGE, pooled, concentrated, and were stored in 10% glycerol at −80 °C for further use. Concentrations for SufS were based on PLP quantification at 388 nm in 0.1 M NaOH (ε_388_ = 6600 M^−1^ cm^−1^).(19) PLP occupancy was determined by comparing the concentration of PLP with the concentation of SufS determined by absorbance at 280 nm (ε_280_ = 49,850 M^−1^ cm^−1^) and was greater than 90% for all enzymes used in this study. The concentration of SufE was determined by absorbance at 280 nm (ε_280_ = 20,970 M^−1^ cm^−1^).

### SufS alanine-NDA assay

SufS cysteine desulfurase activity was measured by monitoring the production of L-alanine using naphthalene-2,3-dicarboxaldehyde (NDA) derivatization through fluorescence detection as developed by Dos Santos et al.(20) Briefly, components of the assay mixture contained 0.25-1 μM SufS (based on PLP absorbance), 50 mM MOPS, pH 8.0, 150 mM NaCl, 2 mM tris(2-carboxyethyl)phosphine) (TCEP) and varying amounts of L-cysteine (0-500 µM) and SufE (0-20 µM). Reactions (50 µL volume) were initiated by the addition of SufS and quenched using 10 μl of 10% trichloroacetic acid (TCA) at various time points ranging from 30 s to 30 min. The quenched reaction mixtures were centrifuged to remove any precipitated protein. Next, 500 μL of freshly prepared NDA-mix containing 100 mM borate, pH 9.0, 2 mM KCN, and 0.2 mM NDA was added to the quenched reaction mixtures and were incubated in the dark for 60 mins. Following incubation, 10 μL of each sample was run on a Zorbax C18 column (Agilent) using a Shimadzu Prominence HPLC with an isocratic gradient of 50% ammonium acetate (pH 6.0) and 50% methanol at the flow rate of 0.4 mL/min. A fluorescence detector was used to detect the Ala-NDA fluorescence adduct (390 nm excitation/440 nm emission). Peak areas were integrated for the Ala-NDA adduct, and the peak area was converted to nanomoles of alanine using an alanine standard curve, made under the same conditions. Initial velocities for alanine formation were determined from at least three time points.

### Fluorescence polarization SufS-SufE binding assay

A fluorescently labeled variant of SufE (C51A/E107C SufE) was obtained as previously discussed.(14) The pET21 plasmid carrying the C51A/E107C SufE gene was transformed into BL21(DE3)*Δsuf E. coli* cells and expressed and purified as described for SufE above with a change in purification buffer pH from 7.5 to 8.0. Following purification and concentration, the protein was dialyzed into 50 mM MOPS (pH 8) and 150 mM NaCl to remove DTT. Fluorescently labeled samples of C51A/E107C SufE were created by incubating 50 µM protein with 500 µM BODIPY FL maleimide (Invitrogen) for 4 h at 25 °C. After the incubation period, unreacted dye was removed using a PD-10 column. Labeling efficiency was determined by comparing the ratio of label concentration (ε_505_ = 80,000 M^−1^cm^−1^) to protein concentration (ε_280_ = 20,970 M^−1^cm^−1^) and was determined to be between 60-80% using this method. SufS-SufE binding titrations were carried out in black 96-well plates in 50 mM MOPS (pH 8), 150 mM NaCl, and 0.1 mg/mL bovine serum albumin. SufS (0.2-80 µM) was mixed with 100 nM labeled C51A/E107C SufE and allowed to equilibrate at room temperature for 30 min. After equilibration, the fluorescence polarization (480 nm excitation, 520 nm emission) was measured using a BioTek Synergy2 multiwell plate reader.

### X- ray structure determination

Crystallization and structure solution of SufS site-directed mutants was performed essentially as previously reported.(14) Briefly, 11-16 mg/mL protein was mixed 1:2 (vol/vol) with crystallization solution, 4-4.5 M NaCl, and 100 mM MES pH 6.5, and subjected to sitting drop vapor diffusion at 20 °C. Crystals appeared after several days. Crystals were cryoprotected by soaking for several minutes in a solution consisting of mother liquor with 50% glycerol. Crystals were plunge-frozen in liquid nitrogen and X-ray data collection proceeded at 100 K with 1.54 Å X-rays generated by a Rigaku Synergy DW equipped with a HyPix6000-HE detector. Data reduction was performed with XDS and merged with XSCALE.(21) The structure given by PDB code 6mr2 (with heteroatoms removed) provided phases for structure solution. Model building and refinement were performed iteratively using Coot and Phenix.(22, 23) X-ray data statistics and model quality data are reported in Table 3. Final coordinates were submitted to the PDB as PDB codes 7rrn (R56A SufS) and 9d2d (R359A SufS).

### Data analysis

Initial velocity data for alanine formation was fit to the Michaelis-Menten equation (Eq 1) to determine the kinetic parameters for each enzyme. Here, *v* is the initial velocity, *E*_t_ is the total enzyme concentration, *k*_cat_ is the maximal turnover number, *S* is the substrate concentration, *K*_M_ is the Michaelis constant. Equation 2 describes substrate inhibtion kinetics with an added inhibition constant (*K*_i_). Fluorescence anisotropy data used to measure protein-protein interactions were fit to equation 3 where *A*_o_ is the anisotropy in the absence of the ligand, *ΔA* is the total change in anisotropy, and *K*_D_ is the dissociation constant.

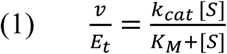

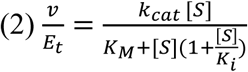

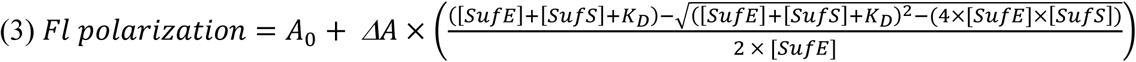

## Results

### Steady-state kinetics suggests that R56 and R359 are required for optimum SufS activity

R56 and R359 SufS residues were separately substituted with either alanine or lysine to generate four SufS variants. The R56 and R359 SufS variant proteins along with WT SufS were expressed and purified to homogeneity as determined by SDS-PAGE (Figure S1). Steady-state kinetic parameters of R56 and R359 SufS variants were evaluated using an HPLC-based fluorescence alanine detection assay.(20) The rate of alanine formation for the SufS enzyme was quantified via NDA-derivatization and HPLC coupled with a fluorescence detector. Kinetic parameters were obtained using the Michaelis-Menten equation. Optimal steady-state activity of SufS requires the presence of both SufE, as a preferred persulfide acceptor, and TCEP, as a preferred reductant for the SufE persulfide. TCEP alone is not sufficient to rapidly reduce the SufS persulfide as the active site is protected from exogenous reductants by the β-latch structural element.(7, 14) Thus, in the presence of TCEP alone, WT SufS has a low SufE-independent activity with a *k_cat_* value of 1.9 ± 0.1 min^−1^ and *K_m_* value of 28 ± 1 μM for cysteine (Table 1, Desulfurase Reaction and Figure 2A). When assayed in the absence of SufE, all four SufS variants exhibit similar *K*_M_ values for cysteine (25-60 µM) suggesting the initial interaction with cysteine is not perturbed. Substitutions with alanine at both positions results in decreases in *k*_cat_ values of 5- and 10-fold for the R56A and R359A variants, respectively. The more conservative R56K substitution is able to fully restore the WT SufS activity, while the R359K substitution partially rescues activity.

**Figure 2.**
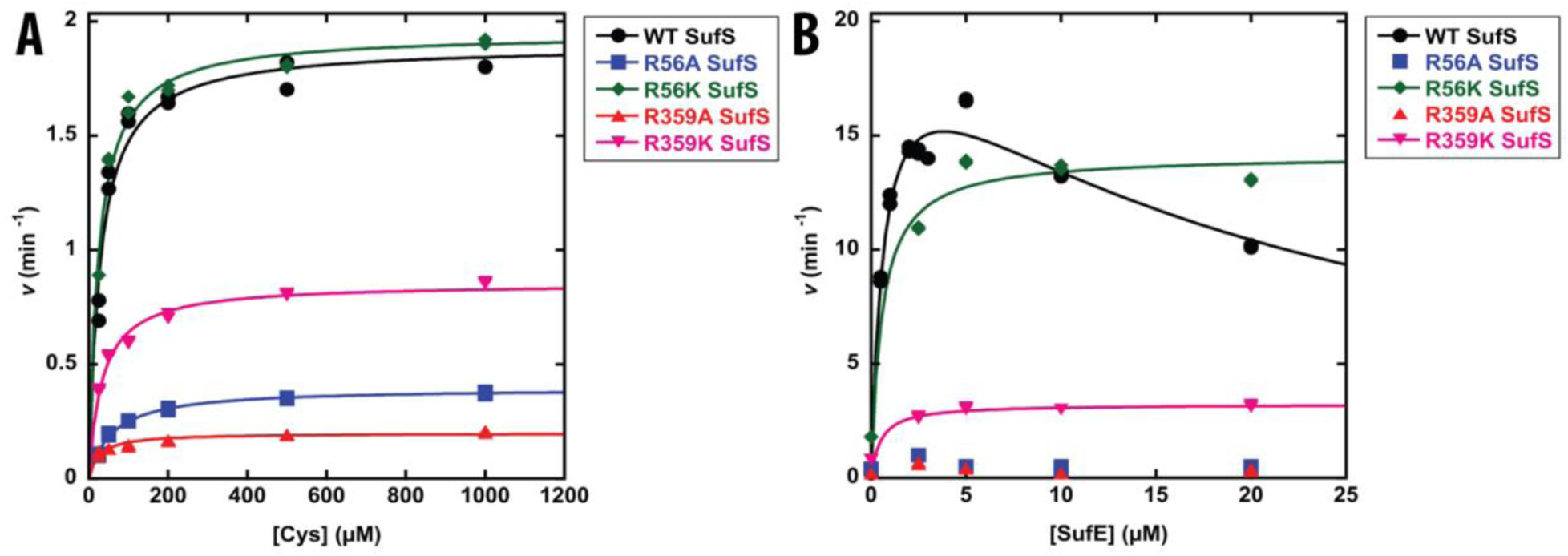
Kinetics of alanine production by WT, R56A, and R359A SufS variants under steady state conditions. Alanine production was measured by HPLC-based assay. (A) Formation of alanine in the absence of SufE. (B) Formation of alanine in the presence of SufE and 500 µM cysteine. Constant reaction conditions for both experiments were 50 mM MOPS pH 8.0, 150 mM NaCl, 2 mM TCEP, and 0.1 to 1.0 μM SufS variant. When SufE was present, SufS concentration was at least 5-fold lower than the SufE concentration in each assay. Solid lines are from fits of the data to equations describing Michaelis-Menten kinetics (equation 1) or substrate inhibition kinetics (equation 2). Duplicate data points are shown but may be obscured by closely overlapping symbols.

**Table 1.** Kinetic parameters for R56 and R359 SufS variants.^a^.

| **Enzyme** | *Desulfurase reaction* |  | *Transpersulfurase reaction* |  |
| --- | --- | --- | --- | --- |
| | $k_{\text{cat}}$ ($\text{min}^{-1}$)<sup>b</sup> | $K_{\text{cys}}$ ($\mu\text{M}$)<sup>b</sup> | $k_{\text{cat(SufE)}}$ ($\text{min}^{-1}$)<sup>c</sup> | $K_{\text{SufE}}$ ($\mu\text{M}$)<sup>c</sup> |
| WT SufS | $1.9 \pm 0.01$ | $28 \pm 1$ | $22 \pm 2^{\text{d}}$ | $0.7 \pm 0.2^{\text{d}}$ |
| R56A SufS | $0.4 \pm 0.01$ | $60 \pm 7$ | not activated<sup>e</sup> | - |
| R56K SufS | $1.9 \pm 0.10$ | $24 \pm 4$ | $13 \pm 1$ | $0.8 \pm 0.3$ |
| R359A SufS | $0.2 \pm 0.01$ | $27 \pm 4$ | not activated<sup>e</sup> | - |
| R359K SufS | $0.9 \pm 0.03$ | $33 \pm 5$ | $2 \pm 0.4$ | $0.5 \pm 1.0$ |

<sup>a</sup>Experiments performed as described in Methods section. <sup>b</sup>Michaelis-Menten kinetic parameters under varying cysteine concentration in the absence of SufE. <sup>c</sup>Michaelis-Menten kinetic parameters under varying cysteine concentration in the presence of SufE. <sup>d</sup>The WT SufS kinetics displays substrate inhibition by SufE contains an addition $K_i$ value of $20 \pm 6 \mu\text{M}$ . Data from reference 8. <sup>e</sup>SufS activity is not improved in the presence of SufE.

In the presence of SufE, the rate of alanine production by SufS is activated ∼10-fold due to the increased rate of persulfide transfer from SufS.(7, 24) Using SufE as a varied substrate under saturating concentrations of cysteine results in substrate inhibition kinetics for WT SufS (Table 1, Transpersulfurase Reaction and Figure 2B).(14) The rate of alanine production by the two alanine substituted variants, R56A and R359A SufS, was not stimulated by the addition of SufE up to 20 µM and remained at the level of the SufE-independent activity (Figure 2B). In contrast, the more conservative lysine substitutions do allow for activation of SufS by SufE and do not appear to be subject to substrate inhibition. Similar to the results above, R56K SufS fully rescued SufE-dependent activity, while the R359K substitution only partially rescued activity. Overall, the conserved arginine residues are important for SufS catalysis with contributions to both alanine formation and persulfide transfer to SufE, with R359 appearing to be more critical for both functions.

### Substitution of R56 or R359 does not negatively perturb SufS/SufE interactions

One explanation for the failure of SufE to activation several of the SufS variants is that the substitution disrupts formation of the SufS/SufE complex. To investigate this possibility, *K*_D_ values for the SufS/SufE complex formation were determined via a fluorescence polarization binding assay.(14) As previously described, 0.05 µM C51A/E107C SufE labeled with BODIPY-FL-maleimide was titrated with the SufS variants (0-20 µM) (Figure 3). Changes in fluorescence polarization were fit to equation 3 to determine a *K*_D_ value (Table 2). Previous studies on WT SufS have shown the presence of cysteine promotes tighter assocation between SufS and SufE with a 10-fold improvement in *K*_D_ value (from ∼5 µM to ∼ 0.5 µM)(14), so binding assays were determined in the absence and presence of 500 µM cysteine. In the absence of cysteine, the R56A, R56K, and R359K variants show *K*_D_ values similar to that determined with WT SufS. In contrast, R359A SufS exhibits a reduced *K*_D_ value in the absence of cysteine suggesting this variant has improved the SufS/SufE interaction. The addition of cysteine causes small decreases in *K*_D_ values for complex formation with the value for R359A remaining lower than the values determined with the other variants. With reasonable *K*_D_ values determined for all SufS variants, the kinetic defects are unlikely to be due to loss of SufE binding.

**Figure 3.**
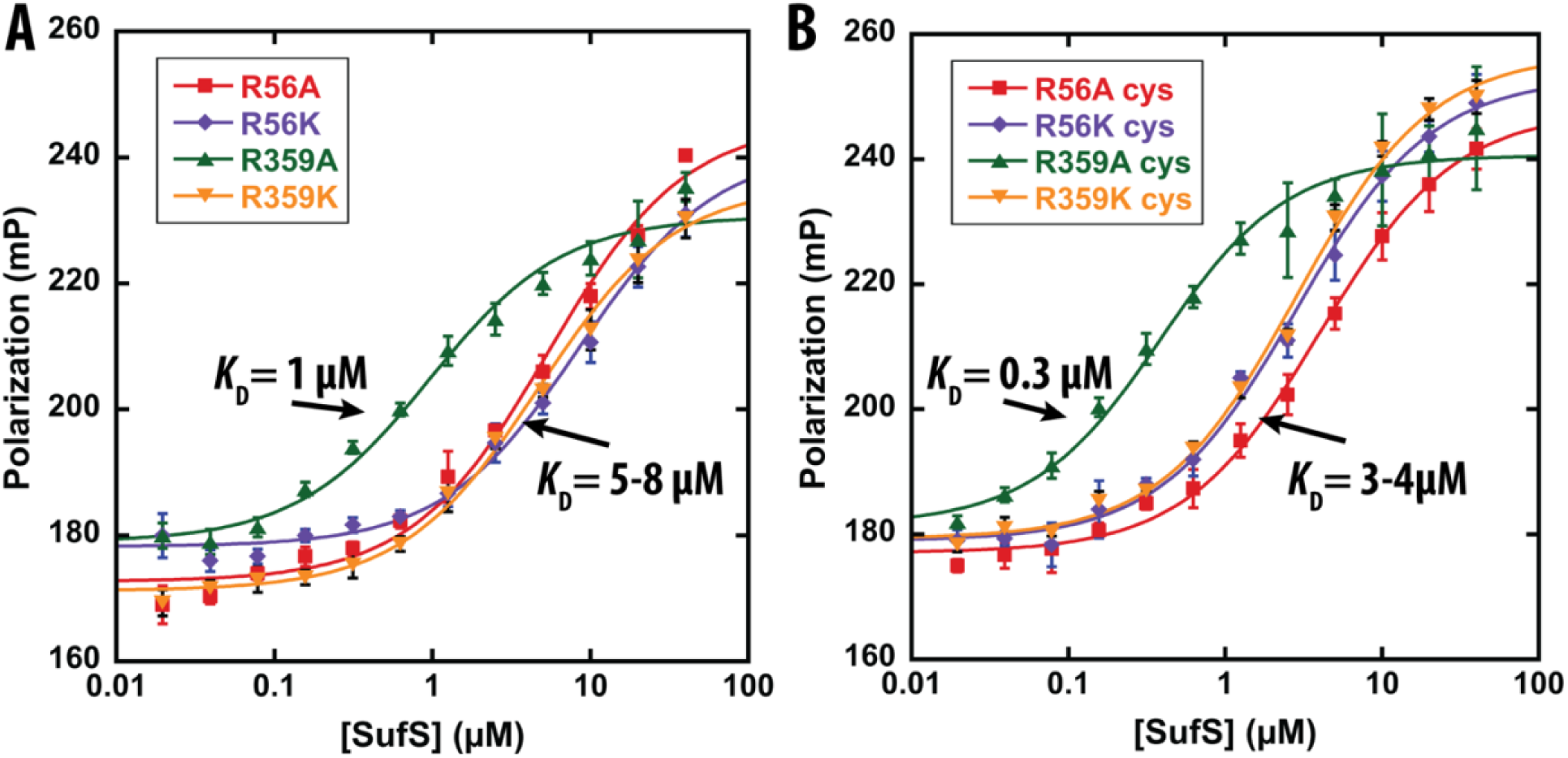
SufS-SufE binding measured by fluorescence polarization. The data show changes in fluorescence polarization due to titration of BODIPY-FL labeled C51A/E107C SufE (0.05 µM) by WT, R56 and R359 SufS variants (0.05 – 20 µM) in the (A) absence and (B) presence of 500 µM cysteine. Error bars are the results of standard deviation from triplicate data. Solid lines in both panels are from a fit of the data to Equation 3. *K*_D_ values determined from the fits are shown in Table 2.

**Table 2.** Dissociation constants for formation of the SufS/SufE complex determined via fluorescence polarization.^a^.

| **Enzyme** | $K_D$ w/o cysteine ($\mu\text{M}$) | $K_D$ w/ cysteine ($\mu\text{M}$) |
| --- | --- | --- |
| WT SufS | $2.5 \pm 0.3$ | $0.2 \pm 0.1$ |
| R56A SufS | $5.8 \pm 0.9$ | $4.0 \pm 0.4$ |
| R56K SufS | $8.5 \pm 1.1$ | $2.8 \pm 0.3$ |
| R359A SufS | $0.9 \pm 0.2$ | $0.3 \pm 0.1$ |
| R359K SufS | $4.6 \pm 0.5$ | $2.8 \pm 0.3$ |

<sup>a</sup>Experiments performed as described in Methods section.

The shift in *K*_D_ values in the presence of cysteine is thought to occur due to formation of a trapped SufS persulfide intermediate that cannot be transferred to the labeled SufE as it lacks the C51 active site residue.(14) This high-affinity SufS/SufE complex has been termed the “close-approach” complex as it likely mimics the conformation leading to persulfide transfer. Thus, comparison of *K*_D_ values for the SufS/SufE complex determined in the presence of cysteine report on both the catalytic fitness of SufS and its ability to form the “close-approach” conformation. Among the R56A, R56K, and R359K variants, none show a decrease in *K*_D_ value on the order of that seen with the WT SufS enzyme. Of these three variants, only R56K SufS exhibits a catalytic ability close to that of the WT enzyme. This suggests that R56K SufS may have trouble forming the “close-approach” complex, while the lack of change in R56A and R359K SufS may derive from low catalytic activity. R359A SufS displayed an unexpected behavior, but one that was rationalized by its crystal structure (below): it shows high-affinity for SufE in the presence and absence of cysteine. As this variant is catalytically inactive, these results suggest that the substitution allows SufS to form a version of the “close-approach” complex in the absence of any catalytic intermediates.

### Both the R56A and R359A substitutions result in conformational changes in the α3-α4 loop on SufS

To better understand the structural reasons for the kinetic defects and altered SufE binding effects of the SufS variants, X-ray crystal structures were determined for the R56A and R359A variants (Table 3). In the R56A SufS structure (PDB ID: 7rrn), residues 56 and 57 are not ordered in the α3-α4 loop (residues 50-59) (Figure 4A). Compared to a high-resolution structure of WT SufS (PDB ID: 6mr2), the adjacent residues of H55 and I58 are displaced slightly from their WT positions. The overall disorder of the loop appears to have minimial effects on positions of the active site residues, and does not provide an immediate explanation to the lack of catalytic activity for the transpersulfurase reaction seen with the R56A variant. Important to discussion later in the text, R359 is found in the out position interacting with the α3-α4 loop.

**Table 3.**
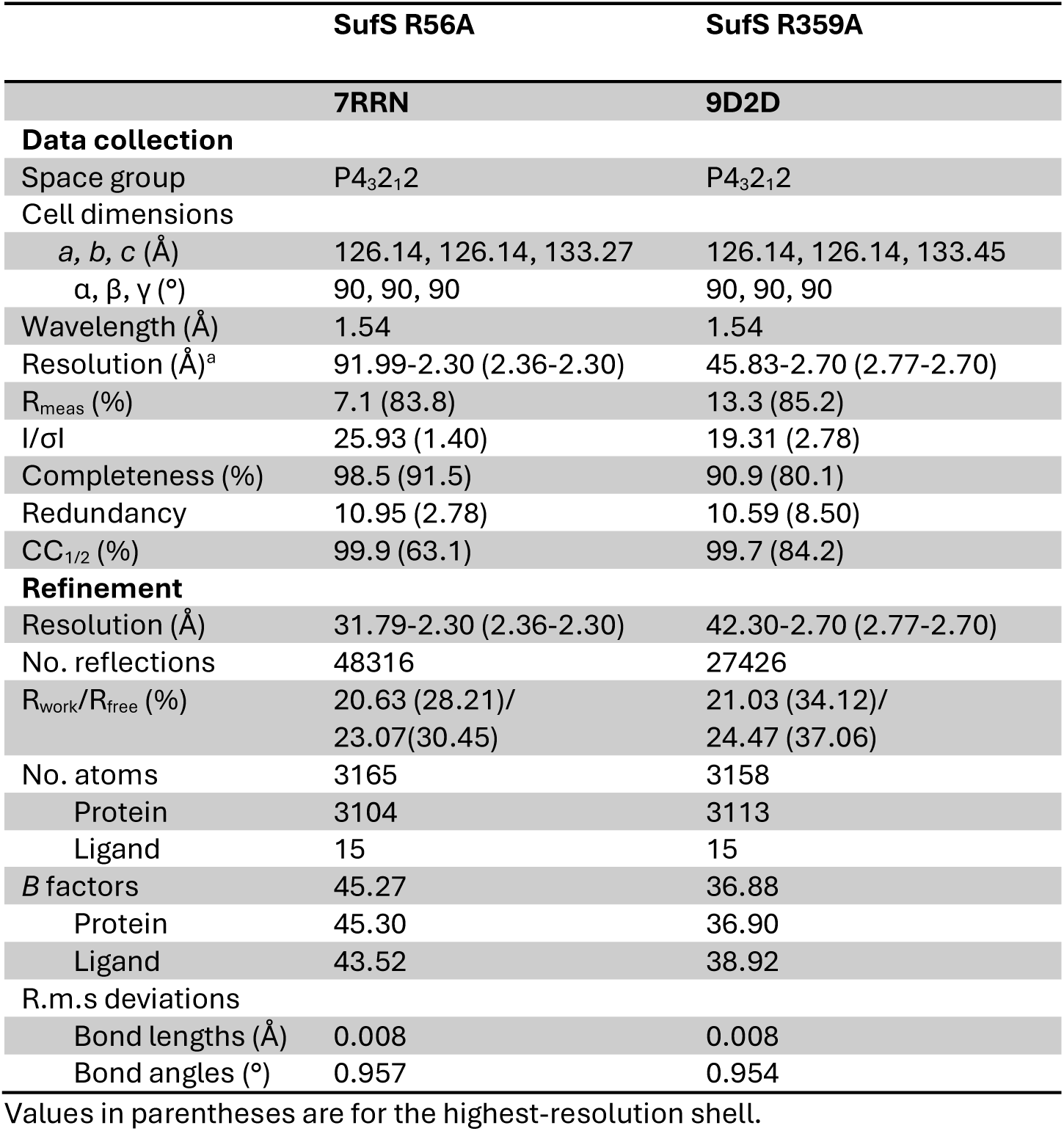
X-ray data statistics.

|  | SufS R56A | SufS R359A |
| --- | --- | --- |
|  | 7RRN | 9D2D |
| **Data collection** |  |  |
| Space group | P4<sub>3</sub>2<sub>1</sub>2 | P4<sub>3</sub>2<sub>1</sub>2 |
| Cell dimensions |  |  |
| *a*, *b*, *c* (Å) | 126.14, 126.14, 133.27 | 126.14, 126.14, 133.45 |
| $\alpha$, $\beta$, $\gamma$ (°) | 90, 90, 90 | 90, 90, 90 |
| Wavelength (Å) | 1.54 | 1.54 |
| Resolution (Å)<sup>a</sup> | 91.99-2.30 (2.36-2.30) | 45.83-2.70 (2.77-2.70) |
| R<sub>meas</sub> (%) | 7.1 (83.8) | 13.3 (85.2) |
| I/σI | 25.93 (1.40) | 19.31 (2.78) |
| Completeness (%) | 98.5 (91.5) | 90.9 (80.1) |
| Redundancy | 10.95 (2.78) | 10.59 (8.50) |
| CC<sub>1/2</sub> (%) | 99.9 (63.1) | 99.7 (84.2) |
| **Refinement** |  |  |
| Resolution (Å) | 31.79-2.30 (2.36-2.30) | 42.30-2.70 (2.77-2.70) |
| No. reflections | 48316 | 27426 |
| R<sub>work</sub>/R<sub>free</sub> (%) | 20.63 (28.21)/<br>23.07(30.45) | 21.03 (34.12)/<br>24.47 (37.06) |
| No. atoms | 3165 | 3158 |
| Protein | 3104 | 3113 |
| Ligand | 15 | 15 |
| *B* factors | 45.27 | 36.88 |
| Protein | 45.30 | 36.90 |
| Ligand | 43.52 | 38.92 |
| R.m.s deviations |  |  |
| Bond lengths (Å) | 0.008 | 0.008 |
| Bond angles (°) | 0.957 | 0.954 |

Values in parentheses are for the highest-resolution shell.

**Figure 4.**
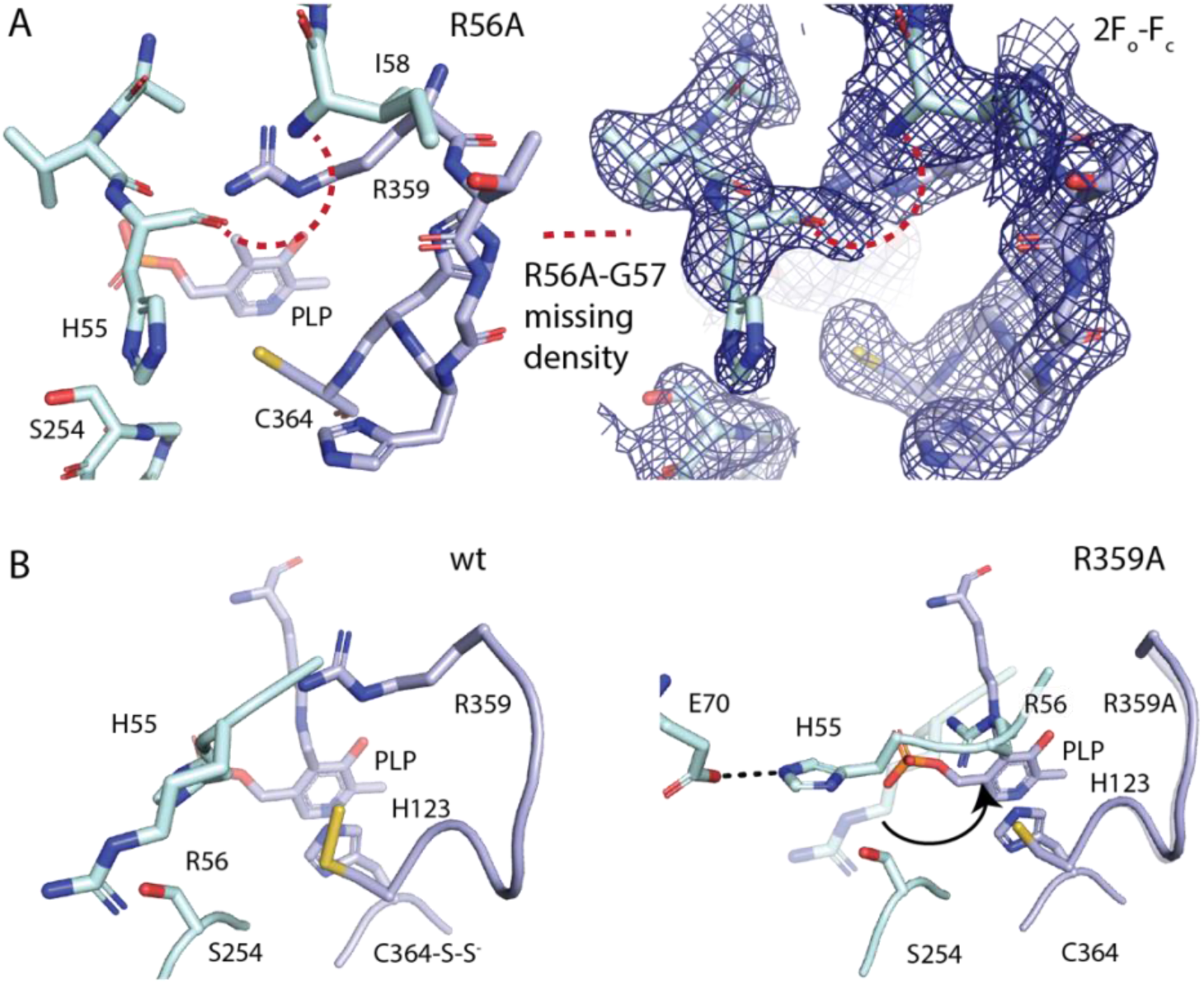
Active site structure of R56A and R359A SufS. (A) The X-ray structure SufS R56A reveals no electron density is present for loop residues R56A and G57, indicating this two amino acid segment has become dynamic in the mutant. R359 is located in the “out” position and interacts with the α3-α4 loop. A 2F_o_-F_c_ electron density map contoured at 1.0σ is shown as blue mesh. (B) A comparison of the X-ray structures of wild type SufS (PDB code 6mr2) versus the R359A mutant reveals that loss of the R359 sidechain promotes a dramatic conformational change in H55, R56 and the α3-α4 loop that now has an inward pointing R56. The conformation with R56 “in” and H55 “out” was previously observed in the PLP-Cys aldimine structure of SufS (PDB cod 6o11). A semi-transparent R56 residue from the wild type structure is shown superpositioned against the R359A mutant to reveal the magnitude (arrow) of the conformational change.

The X-ray crystal structure of R359A SufS also exhibits changes in the α3-α4 loop conformation (Figure 4B). In this structure, H55 and R56 have swapped locations with H55 pointing out of the active site and R56 in the “in” position protruding into the active site. In the inward position, R56 interacts with the sidechain of T278 and is located ∼ 4Å from C364. Superposition of the R359A structure with a C364-persulfide containing structure (PDB ID: 6mr2) shows the R56 sidechain within 3.3 Å of the terminal sulfur atom. In the new position for H55, it interacts with the side chain of nearby E70, located in the adjacent α4 helix. The positioning of the R56 and the α3-α4 loop are in a near identical pose to that seen in the C364A SufS structure that contains a PLP-Cys aldminine intermediate, which originally initiated the interest in these positions. This structural result provides an explanation for the low *K*_D_ values seen with the R359A SufS variant for the SufS/SufE complex. A recent co-crystal of the SufS/SufE complex from *E. coli* shows that positioning of the R56 residue on the α3-α4 loop can physically block the approach of SufE.(17) The R359A-induced “in” conformation of the α3-α4 loop would be expected to result in formation of the “close-approach” SufS/SufE complex in the absence of cysteine, consistent with the binding assay results for this variant.

## Discussion

The SufS cysteine desulfurase from *E. coli* is among the best characterized type II systems. With its dedicated role in Fe-S cluster bioassembly, it is also a model system for the study of protected persulfide transfer in living systems. Recent structures of C364A SufS with a trapped PLP-Cys aldimine and for H123A SufS with a trapped PLP-Cys ketimine intermediate allowed for proposal of specific acid/base roles in catalysis including K226 for removal of the alpha-proton and H123 for deprotonation of C364 prior to nucleophilic attack on the PLP-Cys ketimine intermediate.(16) The C364A SufS structure with a trapped Cys-aldimine intermediate also showed movements in the positioning of H55 and R56 on the α3-α4 loop and residue R359 with both R56 and R359 forming interactions with the sulfur atom of the PLP-Cys aldimine structure. A structure of the SufS/SufE complex from *E. coli* shows that R56 may also play a role in regulating the “close-approach” conformation. To probe the significance of these residues in the SufS desulfurase reaction, site-directed variants of R56 and R359 were generated in SufS and the resulting enzymes were analyzed for effects on catalysis, affinity for SufE, and structure.

### R56 is essential for catalysis and formation of the “close-approach” SufS/SufE complex

H55 and R56 reside on the mobile α3-α4 loop, which undergoes a change in conformation upon formation of the PLP-cys aldimine intermediate with H55 moving from “in” to “out” and R56 moving from “out” to “in”. The SufS H55A variant was previously characterized and showed no detrimental effects on the activity of SufS or the structure.(16) In contrast, the R56A substitution results in loss of catalytic activity. The more conservative substitution of R56K is able to partially restore activity activity suggesting the positive charge of R56 is required. On the basis of *K*_D_ values determined for formation of the SufS/SufE complex, loss of R56 is not detrimental to formation of an initial complex, but it does cause a defect in forming a “close-approach” complex due to the slightly higher *K*_D_ value in the presence of cysteine. The differential response of the H55A and R56A substitutions suggest that although both residues change positions, it is the “in” position of R56 that is of catalytic importance.

### R359 is essential for catalysis and regulating formation of the “close-approach” complex

R359 appears to play a greater role in catalysis compared to R56, based on the result that the R359K still exhibits very poor kinetic parameters. The contribution to catalysis is perhaps not surprising given the location of R359 when it is in the “in” position. The PLP-cys aldimine structure (PDB ID: 6o11) highlights the 3.3 Å proximity of the R359 sidechain to the sulfur atom of the PLP-cys. Superposition of the PLP-cys aldimine SufS structure with a structure containing a C364 persulfide intermediate (PDB ID: 6mr2) shows that R359 in the “in” position would also be placed with 2.5 Å of the SufS persulfide intermediate. Finally, the R359 residue is conserved in other SufS homologs including SufS from *Bacillus subtillis* and *Mycobacterium tuberculosis*. In these organisms, the SufE persulfide acceptor is functionally replaced with SufU. Analysis of the *B. subtillis* and *M. tuberculosis* SufS/SufU structures captured near the point of persulfide transfer shows the equivalent R359 arginine residue (BsSufS R356, MtSufS R368) within 3-4 Å from the incoming SufU cysteine that will act as a persulfide acceptor (Figure S2).(25, 26) In total, structural evidence places the “in” conformation R359 in SufS enzymes at a location where it is able to interact with all facets of persulfide transfer within the system.

One of the more surprising results from this set of experiments is the finding that the R359A SufS variant binds to SufE with high-affinity in the absence of cysteine. Inspection of the structure shows that conformational changes in response to the R359A substitution are limited to changes in the α3-α4 mobile loop. As described above, the new conformation moves R56 from the “out” to the “in” position. A recent structure of the *E. coli* SufS/SufE complex identified R56 as a residue that could physically block SufE from forming a “close-approach” complex.(17) The combined results from the R539A substitution support this hypothesis as the R56 “in” structure correlates with tight binding between R359A SufS and SufE.

### A proposed mechanism of coordinated motion for R359 and R56 in the SufS mechanism

Considering all the data, we propose a new mechanism explaining the coordinated function of R56 and R359 during the SufS catalytic cycle. In the resting state of SufS, R359 is positioned “out” of the active site and interacts with residues and backbone atoms in the α3-α4 mobile loop (Figure 5, left image). This interaction prevents the loop from changing conformation and maintains the “out” position for R56 and the α3-α4 mobile loop. This conformation may allow for interactions between SufS and SufE, but prevents formation of a “close-approach” complex in the absence of a catalytic intermediate in the SufS active site as seen in the recent SufS-SufE structure. Upon formation of catatlyically active intermediates, R359 moves to the “in” position and coordinates persulfide transfer steps (Figure 5, right image). With R359 in the “in” position, the α3-α4 mobile loop is no longer sterically constrained and is allowed to undergo a conformational change to bring R56 to the “in” position and promotes formation of the high-affinity, close-approach complex with SufE. It should be noted that while movement of R359 to the “in” position is required for movement of the α3-α4 mobile loop, it may not be sufficient as the recent SufS/SufE complex structure shows R359 facing “in” with R56 still facing “out”. The identities of catalytic intermediates that trigger the conformational change are not currently clear. The transition may also rely on a conformational change of the β-latch motif, which is also required for formation of the close-approach complex.

**Figure 5.**
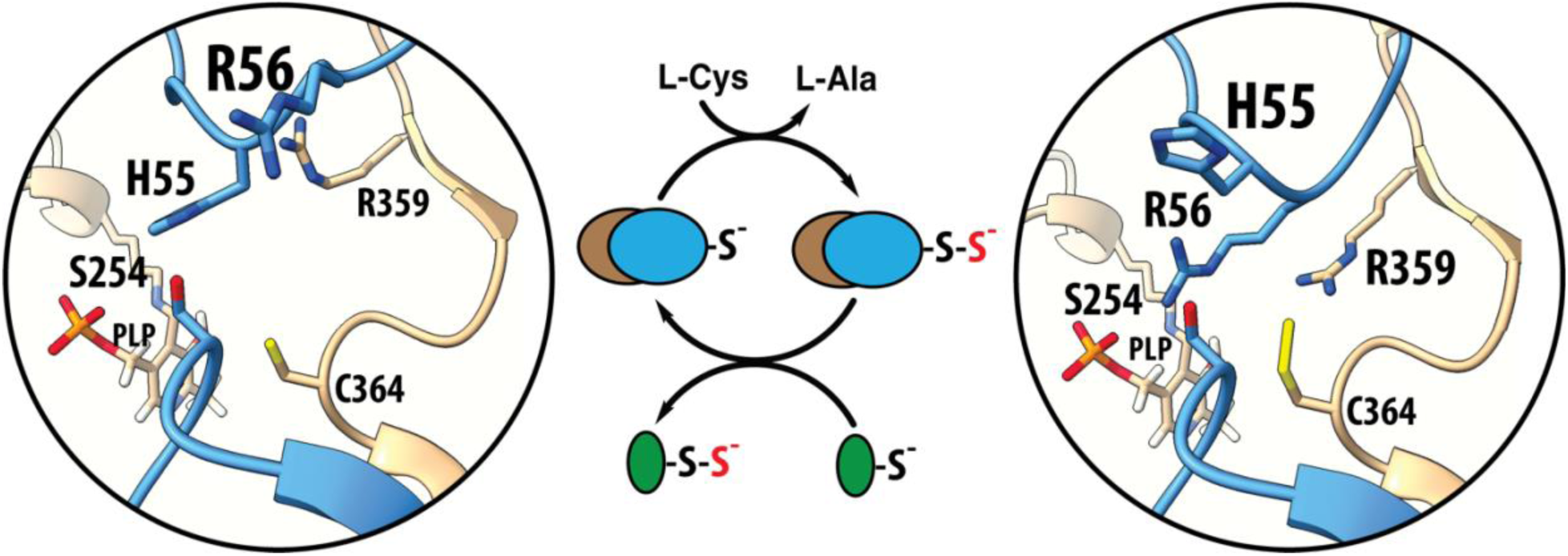
Regulatory model for SufS/SufE complex formation based on movement of the α3-α4 loop. The catalytic cycle of SufS is shown in the center reaction scheme. Models for the resting state (left) and persulfide transfer state (right) of SufS are shown that highlight the coordinated motions of R359 and the α3-α4 loop containing R56.

## Conclusions

In *E. coli,* a PLP-based cysteine desulfurase, SufS, mobilize the sulfur for biosynthesis of iron-sulfur cluster via the SUF pathway.(5) Recent structural, kinetic, and bioinformatic studies of SufS proposed the involvement of residues R56 and R359 in stabilizing PLP-intermediates during SufS catalytic cycle.(16) Here, we have shown that structure and positive charge of the residues at the respective positions is necessary for SufS catalysis. Structures of the two enzyme variants, showed displacement in the α3-α4 mobile loop of SufS, which upon further investigation might also be contributing to the defects observed in SufS-SufE binding. Overall, a coordinated mechanism can be proposed for R56 and R359 SufS residue in response to the changes in the active site, where R359 regulates the positioning of R56 SufS residue.

## Abbreviations

Ala: alanine
Cys: cysteine
DTT: dithiothreitol
K_D_: dissociation constant
MOPS: 3-morpholinopropane-1-sulfonic acid
NDA: naphthalene, 2,3-dicarboxaldehyde
PLP: pyridoxal 5’-phosphate
TCEP: tris (2-carboxyethyl)phosphine

## Acknowledgements

We thank Grace Glidden and Tara Shenkar for their assistance with the initial characterization of the SufS variants.

## References

1. Swindell, J., and Dos Santos, P. C. (2024) Interactions with sulfur acceptors modulate the reactivity of cysteine desulfurases and define their physiological functions Biochim Biophys Acta Mol Cell Res 1871, 119794.

2. Fujishiro, T., Nakamura, R., Kunichika, K., and Takahashi, Y. (2022) Structural diversity of cysteine desulfurases involved in iron-sulfur cluster biosynthesis Biophys Physicobiol 19, 1–18.

3. Baussier, C., Fakroun, S., Aubert, C., Dubrac, S., Mandin, P., Py, B., and Barras, F. (2020) Making iron-sulfur cluster: structure, regulation and evolution of the bacterial ISC system Adv Microb Physiol 76, 1–39.

4. Black, K. A., and Dos Santos, P. C. (2015) Shared-intermediates in the biosynthesis of thio-cofactors: Mechanism and functions of cysteine desulfurases and sulfur acceptors Biochim Biophys Acta 1853, 1470–1480.

5. Blahut, M., Sanchez, E., Fisher, C. E., and Outten, F. W. (2020) Fe-S cluster biogenesis by the bacterial Suf pathway Biochim Biophys Acta Mol Cell Res 1867, 118829.

6. Dussouchaud, M., Barras, F., and Ollagnier de Choudens, S. (2024) Fe-S biogenesis by SMS and SUF pathways: A focus on the assembly step Biochim Biophys Acta Mol Cell Res 1871, 119772.

7. Selbach, B. P., Pradhan, P. K., and Dos Santos, P. C. (2013) Protected sulfur transfer reactions by the *Escherichia coli* Suf system Biochemistry 52, 4089–4096.

8. Dai, Y., and Outten, F. W. (2012) The *E. coli* SufS-SufE sulfur transfer system is more resistant to oxidative stress than IscS-IscU FEBS Lett 586, 4016–4022.

9. Outten, F. W., Wood, M. J., Munoz, F. M., and Storz, G. (2003) The SufE protein and the SufBCD complex enhance SufS cysteine desulfurase activity as part of a sulfur transfer pathway for Fe-S cluster assembly in *Escherichia coli* J Biol Chem 278, 45713–45719.

10. Ollagnier-de-Choudens, S., Lascoux, D., Loiseau, L., Barras, F., Forest, E., and Fontecave, M. (2003) Mechanistic studies of the SufS-SufE cysteine desulfurase: evidence for sulfur transfer from SufS to SufE FEBS Lett 555, 263–267.

11. Cupp-Vickery, J. R., Urbina, H., and Vickery, L. E. (2003) Crystal structure of IscS, a cysteine desulfurase from *Escherichia coli* J Mol Biol 330, 1049–1059.

12. Smith, A. D., Agar, J. N., Johnson, K. A., Frazzon, J., Amster, I. J., Dean, D. R., and Johnson, M. K. (2001) Sulfur transfer from IscS to IscU: the first step in iron-sulfur cluster biosynthesis J Am Chem Soc 123, 11103–11104.

13. Lima, C. D. (2002) Analysis of the *E. coli* NifS CsdB protein at 2.0 A reveals the structural basis for perselenide and persulfide intermediate formation J Mol Biol 315, 1199–1208.

14. Gogar, R. K., Carroll, F., Conte, J. V., Nasef, M., Dunkle, J. A., and Frantom, P. A. (2023) The beta-latch structural element of the SufS cysteine desulfurase mediates active site accessibility and SufE transpersulfurase positioning J Biol Chem 299, 102966.

15. Dunkle, J. A., Bruno, M. R., and Frantom, P. A. (2020) Structural evidence for a latch mechanism regulating access to the active site of SufS-family cysteine desulfurases Acta Crystallogr D Struct Biol 76, 291–301.

16. Blahut, M., Wise, C. E., Bruno, M. R., Dong, G., Makris, T. M., Frantom, P. A., Dunkle, J. A., and Outten, F. W. (2019) Direct observation of intermediates in the SufS cysteine desulfurase reaction reveals functional roles of conserved active-site residues J Biol Chem 294, 12444–12458.

17. Gogar, R. K., Chhikara, N., Vo, M., Gilbert, N. C., Dunkle, J. A., and Frantom, P. A. (2024) The structure of the SufS-SufE complex reveals interactions driving protected persulfide transfer in iron-sulfur cluster biogenesis J Biol Chem 300, 107641.

18. Kim, D., Singh, H., Dai, Y., Dong, G., Busenlehner, L. S., Outten, F. W., and Frantom, P. A. (2018) Changes in Protein Dynamics in *Escherichia coli* SufS Reveal a Possible Conserved Regulatory Mechanism in Type II Cysteine Desulfurase Systems Biochemistry 57, 5210–5217.

19. Peterson, E. A., and Sober, H. A. (1954) Preparation of Crystalline Phosphorylated Derivatives of Vitamin B6 J Am Chem Soc 76, 169–175.

20. Addo, M. A., Edwards, A. M., and Dos Santos, P. C. (2021) Methods to Investigate the Kinetic Profile of Cysteine Desulfurases Methods Mol Biol 2353, 173–189.

21. Kabsch, W. (2010) Xds Acta Crystallogr D Biol Crystallogr 66, 125–132.

22. Liebschner, D., Afonine, P. V., Baker, M. L., Bunkoczi, G., Chen, V. B., Croll, T. I., Hintze, B., Hung, L. W., Jain, S., McCoy, A. J., Moriarty, N. W., Oeffner, R. D., Poon, B. K., Prisant, M. G., Read, R. J., Richardson, J. S., Richardson, D. C., Sammito, M. D., Sobolev, O. V., Stockwell, D. H., Terwilliger, T. C., Urzhumtsev, A. G., Videau, L. L., Williams, C. J., and Adams, P. D. (2019) Macromolecular structure determination using X-rays, neutrons and electrons: recent developments in Phenix Acta Crystallogr D Struct Biol 75, 861–877.

23. Emsley, P., Lohkamp, B., Scott, W. G., and Cowtan, K. (2010) Features and development of Coot Acta Crystallogr D Biol Crystallogr 66, 486–501.

24. Gogar, R. K., and Frantom, P. A. (2024) Persulfide Transfer to SufE Activates the Half-Sites Reactivity of the *E. coli* Cysteine Desulfurase SufS Biochemistry 63, 1569–1577.

25. Fujishiro, T., Terahata, T., Kunichika, K., Yokoyama, N., Maruyama, C., Asai, K., and Takahashi, Y. (2017) Zinc-Ligand Swapping Mediated Complex Formation and Sulfur Transfer between SufS and SufU for Iron-Sulfur Cluster Biogenesis in *Bacillus subtilis* J Am Chem Soc 139, 18464–18467.

26. Elchennawi, I., Carpentier, P., Caux, C., Ponge, M., and Ollagnier de Choudens, S. (2023) Structural and Biochemical Characterization of *Mycobacterium tuberculosis* Zinc SufU-SufS Complex Biomolecules 13, 732.

