## Supporting Information for "Two conserved arginine residues facilitate C-S bond cleavage and persulfide transfer in Suf family cysteine desulfurases"

Table S1. Oligonucleotide primers used in this study

| Variant | Primer | Primer Sequence (5’-3’) |
|---|---|---|
| R56A SufS | Forward | GCGGTGCATGCGGGTATTCATACC |
| | Reverse | GGTATGAATACCCGCATGCACCC |
| R56K SufS | Forward | GCGGCGGTGCATAAAGGTATTCATACC |
| | Reverse | GGTATGAATACCTTTATGCACCGCCGC |
| R359A SufS | Forward | ATTGCTGTGGCGACCGGACATCAC |
| | Reverse | GTGATGTCCGGTCGCCACAGCAAT |
| R359K SufS | Forward | GGCATTGCTGTGAAAACCGGACATCAC |
| | Reverse | GTGATGTCCGGTTTTCACAGCAATGCC |

**Figure S1. SDS-PAGE analysis of purified WT and variant SufS enzymes.**

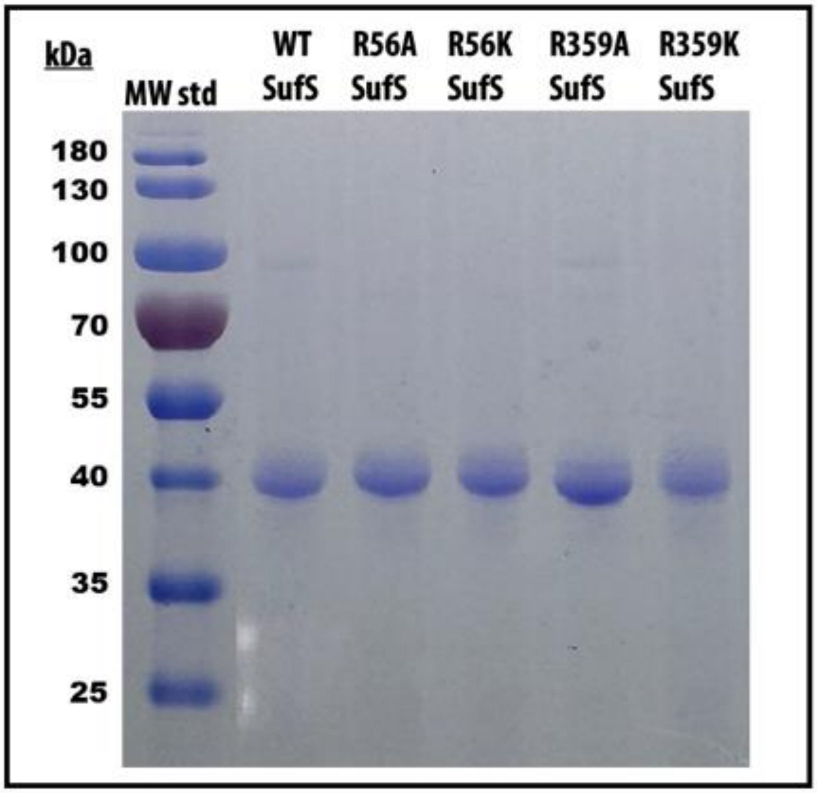

**Figure S2. Positions of R56- and R359-equivalent residues in SufS/SufU co-crystal structures.** Active sites of SufS/SufU from (A) *B. subtilis* (PDB ID: 5xt6) and (B) *M. tuberculosis* (PDB ID:8odq) with labels for key residues.

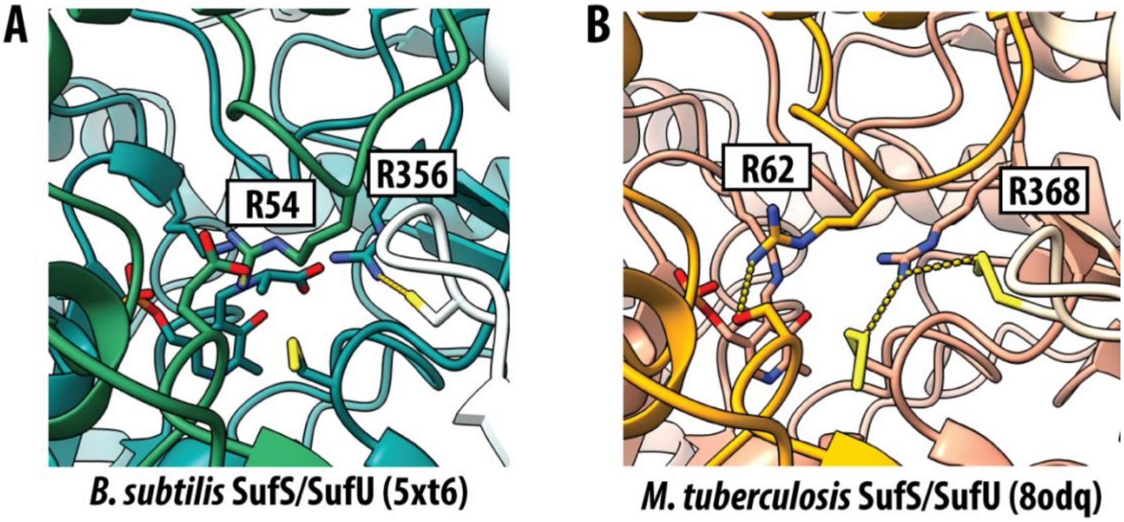
